# A Computational Modeling Framework to Analyze Synovial-Tissue Based Drug Targets and Diagnostic Biomarkers in Rheumatoid Arthritis

**DOI:** 10.1101/2021.11.08.467813

**Authors:** Paridhi Latawa, Brianna Chrisman

## Abstract

Rheumatoid Arthritis (RA) is an inflammatory autoimmune disease that affects 23 million people worldwide. It is a clinically heterogeneous disorder characterized by the attack of inflammatory chemicals on the synovial tissue that lines joints. It is advantageous to develop effective, targeted treatments and identify specific diagnostic biomarkers for RA before extensive joint degradation, bone erosion, and cartilage destruction. Current modes of RA treatments have alleviated and notably halted the progression of RA. Despite this, not many patients reach low disease activity status after treatment, and a significant number of patients fail to respond to medication due to drug non-specificity. While the reasons for these rates remain unknown, the cellular and molecular signatures present in the synovial tissue for RA patients likely play a role in the varied treatment response. Thus, a drug that particularly targets specific genes and networks may have a significant effect in halting the progression of RA. This study evaluates and proposes potential drug targets through *in silico* mathematical modeling of various pathways of interest in RA. To understand how drugs interact with genes, we built a mathematical model with 30 two-gene and three-gene network interactions and analyzed the effect of 92 different perturbations to rate constants. We determined that inhibition of the LCK-CD4, VAV1-CD4, and MLT-ROR pathways could potentially serve as drug targets. We also found that increased activity of the DEC2-IL1β and the NF-kB-interleukin pathway and the decreased activity of the TNF-α-REV-ERB pathway could serve as diagnostic biomarkers.

## Introduction

Rheumatoid Arthritis (RA) is an autoimmune disease affecting more than 23 million people (1, 2). It is characterized by the attack of inflammatory chemicals on the synovial tissue lining joints in the hands, knees, and wrist. The lining of joints becomes inflamed, which can cause damage to joint tissue. Uncontrolled inflammation can lead to cartilage destruction, joint deformities, and painful bone erosion. RA is a systemic disease, meaning it has a cascading effect in terms of the overall deterioration of health (3). With increased inflammation comes a greater probability of developing other complications such as atherosclerosis, anemia, and pericarditis (3). Adults with RA are also less likely to be employed as opposed to those without RA. A recent study found that after five years of onset, approximately 30% of patients had quit their jobs due to RA (4). With common effects of RA being chronic pain, unsteadiness, deformity, and a lack of general everyday function, it becomes vital to provide adequate treatment options for patients.

It is advantageous to develop treatments for RA before extensive erosion sets in, as treatments are more effective at the early stages (5). Current varying modes of treatments for RA include medication known as nonsteroidal anti-inflammatory drugs (NSAIDs), conventional and targeted synthetic disease-modifying antirheumatic drugs (DMARDs), steroids, and biologic agents (6). Although several treatments have been found to alleviate and notably halt the progression of RA, a significant number of patients still fail to successfully and fully respond to current medication due to the non-specificity of the drugs (7). As measured by the American College of Rheumatology (ACR) response criteria, only 20-30% of patients reach low disease activity status after treatments. While the mechanistic reason for such failure rates remains unknown, the cellular and molecular signatures of the synovial tissue for RA patients are likely to play a role in the variable treatment response and heterogeneous clinical evolution.

Some of the current medicines that treat RA include Humira, Plaquenil, and Methotrexate (Table 1). Although these medications are effective to a certain extent, they include severe side effects, and a one drug fits all approach often does not have high rates of effectiveness in inhibiting RA (8). It is suggested that patients try varying drug products to determine which one is most efficient, yet this approach can have a low cost-benefit payoff. Methotrexate, in particular, is a drug that can cause severe and life-threatening side effects and should only be taken in severe conditions (9). If medication is not effective, surgeries such as synovectomies, tendon repair, joint fusion, and total joint replacement can be used to improve function and alleviate pain (5). Yet, surgeries also carry a prevalent risk of bleeding, infection, and pain, in addition to being costly and time-consuming. Overall, various drugs are not safe and effective for treating RA (10). Thus, a drug targeted for a particular gene and network may have a significant effect on the current patient and in stopping the genetic transmissions of disease-causing agents in Rheumatoid Arthritis.

**Table 1:** Medication types used to treat Rheumatoid Arthritis (6).

| Medication Type | Drug Examples | Function | Side Effects |
| --- | --- | --- | --- |
| **NSAIDs** | Ibuprofen (Advil, Motrin IB), naproxen sodium (Aleve), etc. | Reduce inflammation and pain | Stomach irritation, heart problems, kidney damage |
| **DMARDs** | Methotrexate, Plaquenil, Azulfidine | Slow progression of RA. Save joints & tissues from permanent damage | Vary - liver damage, severe lung infection |
| **Targeted synthetic DMARDs** | Rinvoq, Olumiant | Used if regular DMARDs and biologics aren't effective. | High doses can increase the risk of blood clots in lungs, heart-related events, cancer |
| **Steroids** | Corticosteroid Medications (prednisone) | Reduction of inflammation, pain, and joint damage | Bone thinning, weight gain, diabetes |
| **Biological Agents** | Humira, Cimzia, Simponi, Rituxan, Actemra | Generally paired with a DMARD like methotrexate. A newer class of DMARDs | Increased risk of infection |

Notably, the inflamed synovial tissue has been found to have great potential for passive and active drug targeting due to its enhanced permeability and retention effect (10). This tissue also consists of numerous RA synovial macrophages and fibroblasts that express surface receptors. Inflammation of joints can lead to intensive joint degradation, as when inflammation occurs in the subchondral bone, it can cause inflammatory cysts and erosion (11). Synovitis, or increased inflammation of the synovium, has been identified as a precursor of erosion. Drugs such as Tocilizumab (Amectra), when treated in tandem with methotrexate and tumor necrosis factor inhibitors (TNF-i), have been developed to interfere in joint destruction by inhibiting IL-6 (12). This study aims to similarly identify drug targets to alleviate joint destruction and eventually inflammation to halt RA’s progression.

In addition to identifying possible drug targets, it is imperative to identify more accurate early diagnostic biomarkers for RA. Early detection has minimized joint destruction and improved patients’ treatment outcomes (13). While ACR and EULAR criteria are based primarily on blood tests to measure serologic risk factors for RA, they do not necessarily reflect the biological actions occurring in the target synovial tissue of the patient (14). Thus, comes the need to identify more specific early diagnostic biomarkers for RA.

To evaluate potential drug targets and diagnostic biomarkers, we mathematically modeled various pathways of interest in RA *in silico*. Biological computational modeling is an important subfield in systems biology that is vital to analyze interactions in biological systems for better disease detection, diagnosis, and treatment. Mathematical models are useful in biology to bridge the understanding of proposed molecular interactions and their resultant tissue-level effects (15). They save lab and testing time by providing more accurate results and data from which proposed avenues of interest can be explored further. Differential equation-based models are a rising development in biology as they take discrete-time and state measurements (16). Many of the best computational models developed for biological processes are expressed through sets of ordinary differential equations (ODEs). ODEs represent how individual variables or connections evolve over time. ODEs often encode a simplified version of a biological system through modeling. These models are based on the premise that the system is well-mixed, meaning it contains a sufficient variety of components such as small molecules, cells, and genes so their numbers can be considered continuous (17).

## Methods

### Software and Libraries

Google Colaboratory (Colab) was used to build the biological model of the pathways of interest for the analysis of RA pathways, and Python was used to program the model (18). The numpy library, matplotlib.pyplot functions, networkx, and seaborn packages as well as the pandas library were imported for the creation of graphs, figures, and tables. For labeling, label lines were also imported. In addition to a cumulative graph of the baseline RA and perturbation model, various heat maps were created for each quantity to measure the differences in final expression levels or concentrations.

### Ordinary Differential Equations (ODEs)

Ordinary differential equations were constructed for each pathway interaction. These equations had a general form based on fundamental properties highlighted by Covert and Liu et al (19, 20).

All the differential equations used have a general form represented by the following (19).

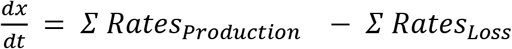

This equation relates the change in amount or concentration of a quantity over time to the sum of the rates of production minus the sum of the rate of loss.

As most of the interactions studied were between gene transcripts, as opposed to proteins, the interactions were derived such that expression was measured and mathematically depicted.

The base equation for messenger ribonucleic acid (mRNA) transcripts was represented as:

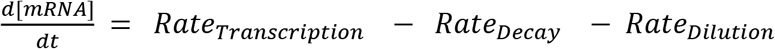

Various main assumptions that were made to simplify this ODE were the following. Assuming a given rate is negligible with respect to the other term in the equation, it is feasible to simplify this term as appropriate. In mRNA, loss by decay is usually faster than the dilution rate, so we can approximate the dilution rate as 0 (19). In small molecules, loss by dilution is generally quicker than the decay/degradation rate, so we can assume the decay rate to be 0 (19). Secondly, we assume that a rate constant for the system does not change significantly under the present condition. Thirdly, we assumed that for small molecules and transcripts, the process follows mass action kinetics such that the reaction rate is proportional to the product of the reactant concentrations. This means that mRNA decay and degradation depend on a reaction between mRNA molecules and ribonuclease enzymes. These relationships are again simplified by defining a kinetic constant to represent relationships. By applying these assumptions, the base equation for mRNA transcription is adjusted to the following where *f*(*G*_*i*_[*t*])represents a function of the different genes that are influencing the rate of mRNA transcription.

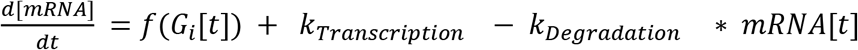

The equations provided by Liu et al were explicitly for proteins and where they adjusted to reset mRNA expression over time (20). From base equations for mRNA activator and inhibitor relationships, we derived equations specific to each interaction.

The base activator equation was derived based on the interactions between three genes: G1, G2, G3. G1 represents an initial gene that is directly activating G2, G2 is a gene directly activating G3, and G3 is a gene that is directly inhibiting G1.

The base activator equation for G2, for example, would be represented as the following.

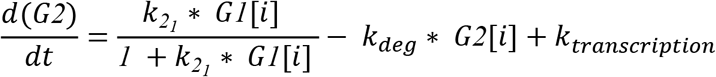

The 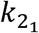 term represents the constant of the rate of change of G1 affecting G2, k_deg_ is the degradation constant that is the same for all terms, and k_transcription_ is the transcription constant for each of the mRNA transcripts. Time is taken in increments, and the overall rate of change is measured. Derivation for constants is explained further in the next section.

The base inhibitor equation was derived similarly as above. For G1, the base inhibitor equation would be the following.

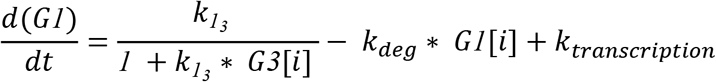

The base equation for small molecules was represented as the following, where k_i_ represents the constant between interactors.

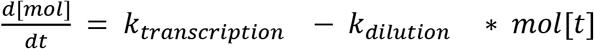

The base equation for small molecules was developed as the following where *f*(*G*_*i*_[*t*]) represents a function of a variety of different genes or components that are influencing the rate of the small molecules.

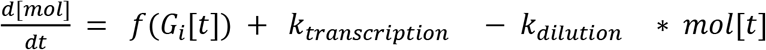

The above base equations were used to derive equations for each of the respective interactions.

The combined differential equations for each interaction are shown in Table 2.

**Table 2:**
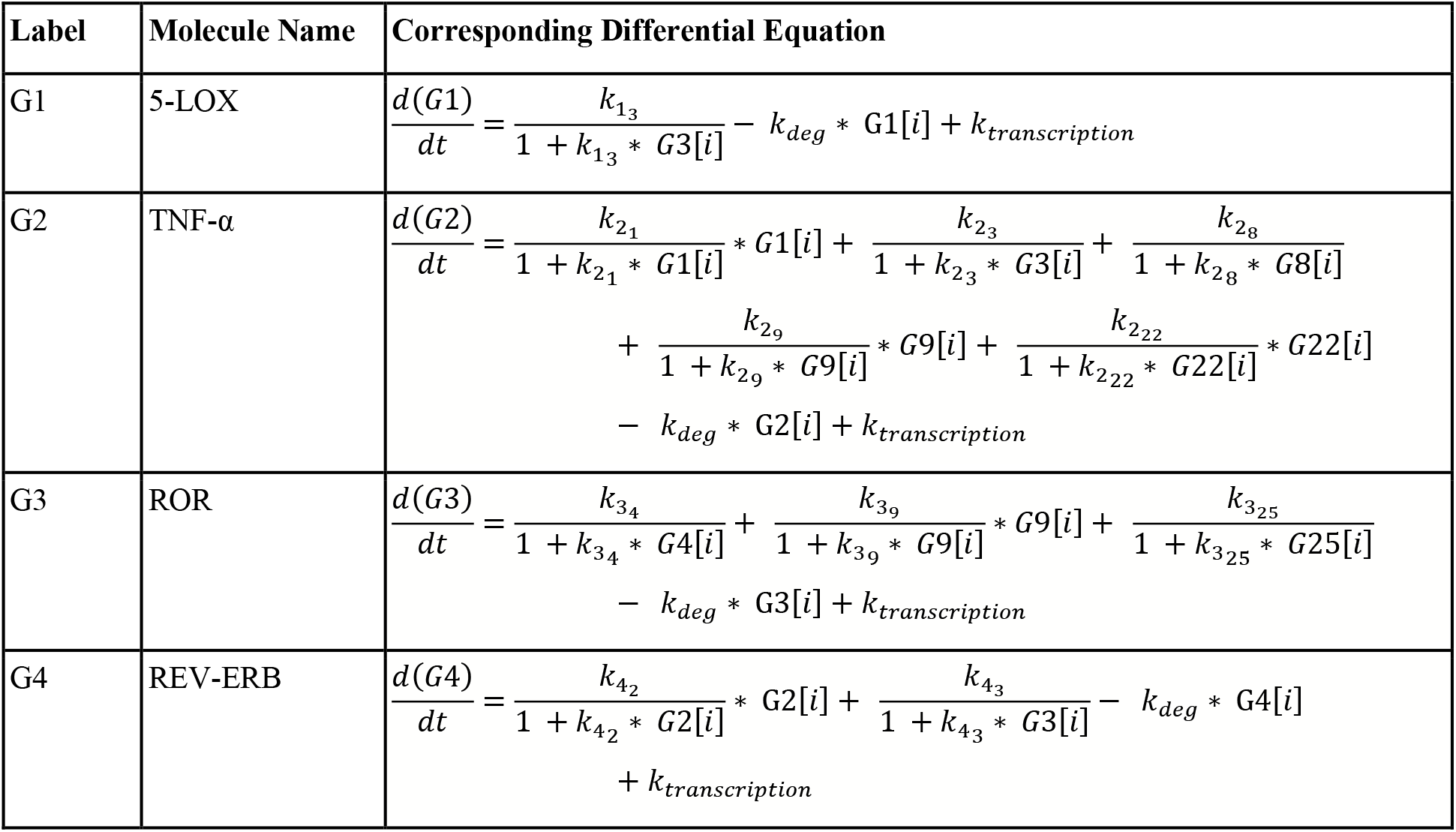
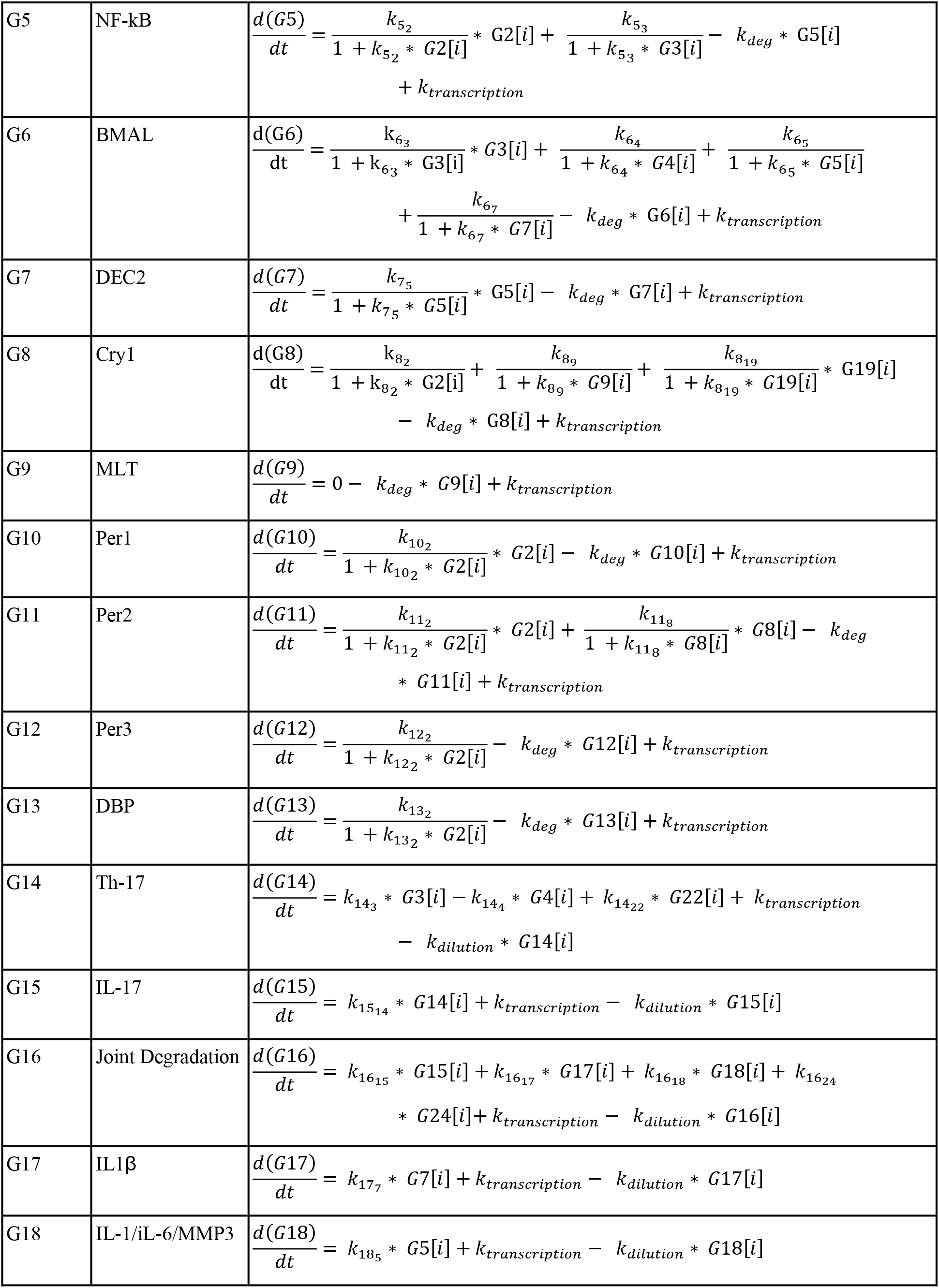
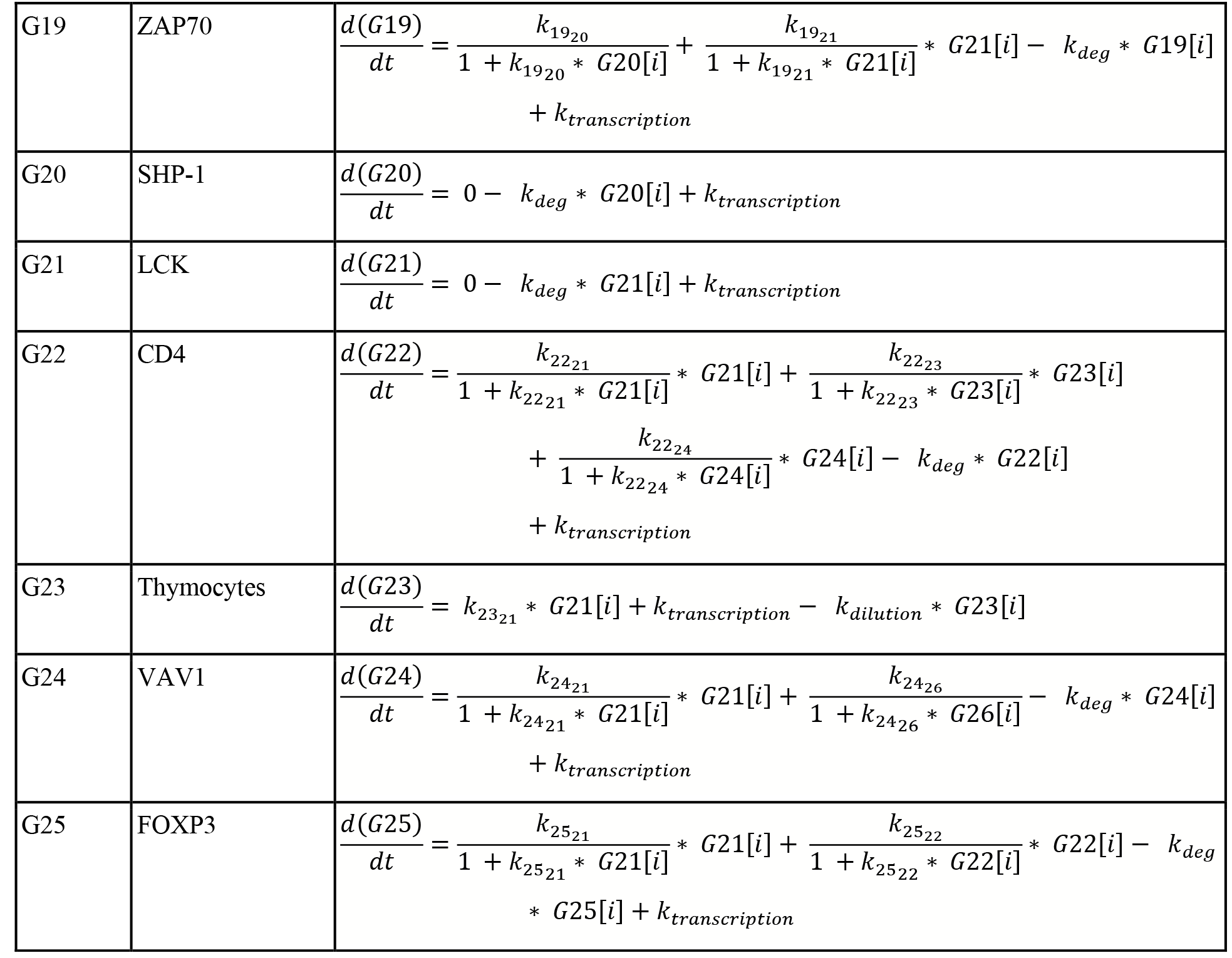
Combined differential equation for each molecule, gene, or process. The leftmost column gives the assigned label & middle column provides the name of the molecule, gene, or process. The right hand column gives the corresponding combined differential equation respective to the molecule, gene, or process.

### Constants and Concentrations

The main constants used were the degradation constant, transcription constant, dilution constant, and the interaction constants between genes. The values of the constants were determined from the literature.

A degradation constant k_deg_ representing the degradation rate of each transcript or gene was incorporated in each of the differential equations. The optimal value for the degradation constant k_deg_ was set to 0.002 sec^−1^, to represent its relationship as a quantity that occurs approximately one magnitude slower than transcription. This degradation rate was a first-order constant and was multiplied by the concentration of the gene transcript at specific intervals in time.

A dilution constant k_dilution_ was incorporated in the differential equations for small molecules or cells. This constant represents how small molecules could get used in unrelated pathways that are not used in the RA reaction or diffuse out of the cell such that they are not easily reachable. For example, interleukins are used in a variety of pathways in addition to the RA pathway, so dilution constants are needed to represent this. The dilution constant was a first-order constant and was multiplied by the concentration of the small model or cells at specific intervals of time. No dilution constant was incorporated for genes because, in mRNA, degradation overpowers dilution (19).

Assuming that most of the proteins are being transcribed when they do not have an activator bound, we also incorporated a transcription constant k_transcription_ representing the baseline transcription rate. Bio Numbers states that a one kb gene should take at maximal transcription rate about 1000 nt/80 nt/s ≈ 10s,” (21). A human gene is usually ~10kb long, so the transcription rate of a gene transcription = 0.01 sec^−1^ was used in the model. This transcription rate was zeroth order and represented by a single addition in each equation.

Each interaction was also assigned a constant. The constants were written in the format k_α_β_ where α is the model whose rate of change is being examined and β is the molecule who is affecting the rate of change of molecule α. These constants were all assigned a theoretical value of 1 and were modified when incorporating perturbations. We simulated this system and measured the change in concentrations for each type of transcript and small molecule over time.

### Baseline Rheumatoid Arthritis Simulation

The baseline RA model simulating elevated conditions was developed for 26 different molecules and 30 total groups of interactions. All molecules were shown to interact and eventually play a role in joint degradation. Differential equations for individual interaction were first developed individually for each interaction and then combined into a full equation for each molecule.

The base model for interactions in RA was taken from a study done by Jahanban-Esfahlan et al (30). Further pathways of interest were added based on findings of relationships and interactions in literature. More interactions of interest were incorporated based on findings from the literature. The final model was developed in slides and exported as shown in Figure 2.

**Figure 1:**
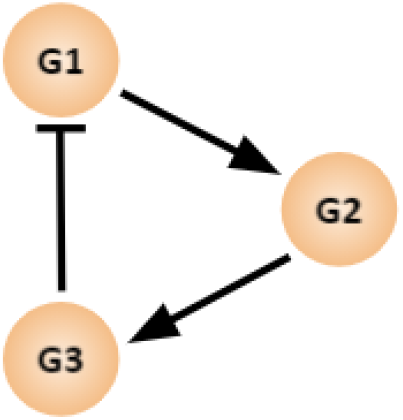
Three gene node networks depicting G1, G2, G3, for which the example equations are derived below.

**Figure 2:**
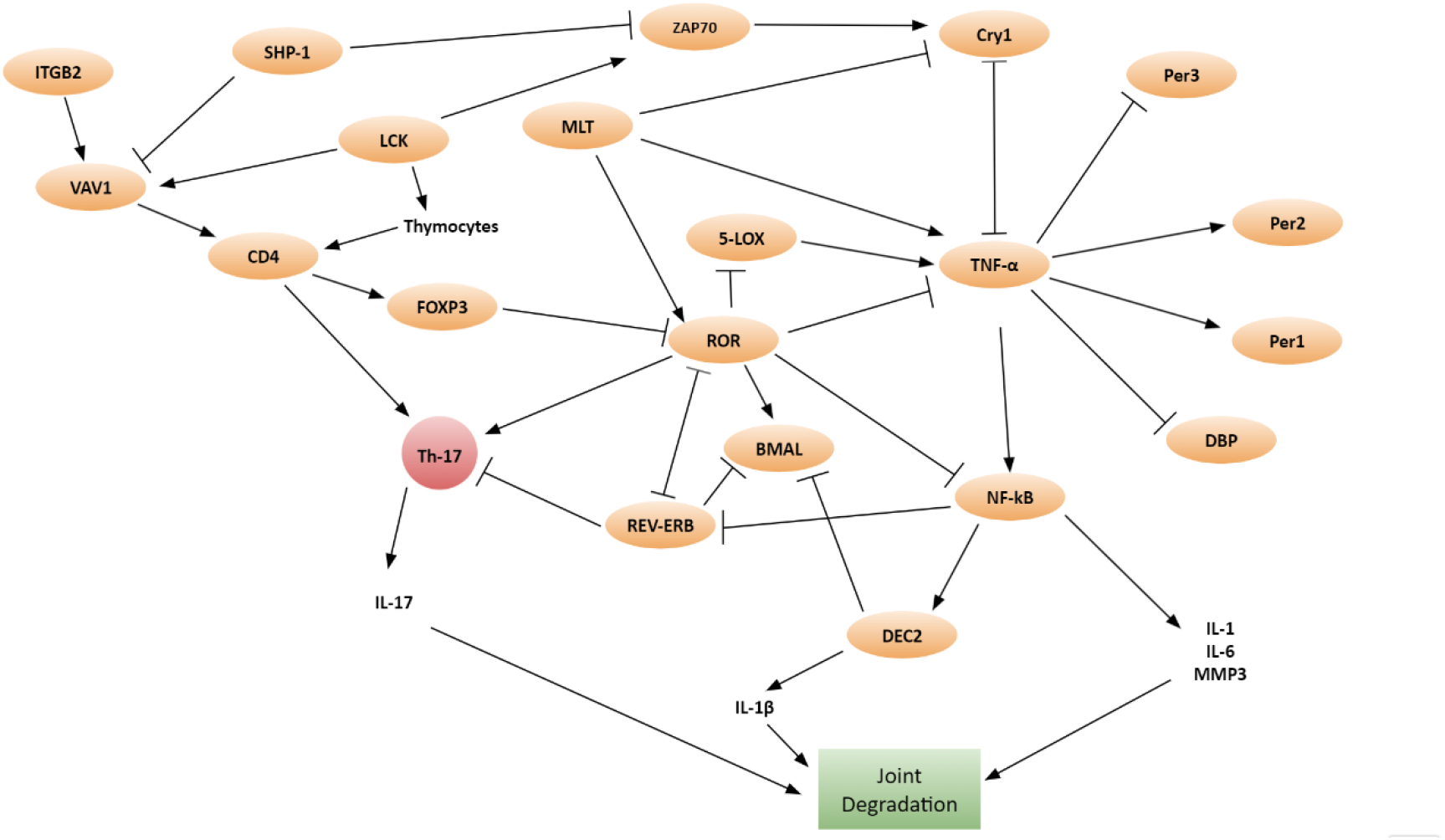
Adjusted RA model with added interactions. An arrow represents activation and a “T” represents inhibition. Orange circles represent genes while components in no boxes or in red represent small molecules.

Nine pathways of interest were added to the original model to create the adjusted RA model based on findings in the literature, as shown in Table 3. 30 total two-gene and three-gene network interactions were modeled.

**Table 3:**
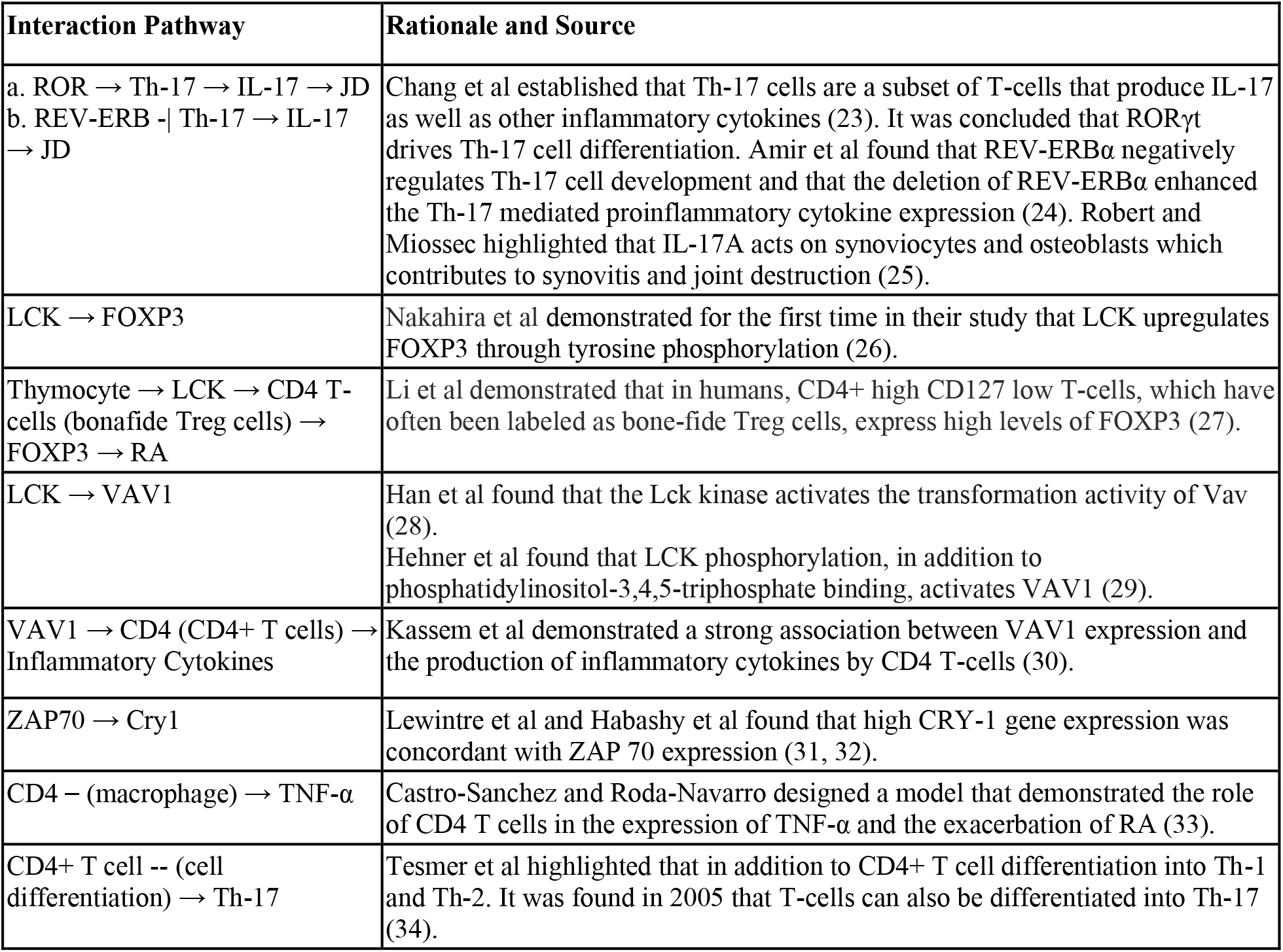
Table presenting rationales for the pathways added, where JD represents joint degradation.

30 total interactions composed of three-gene networks or two-gene pathways were modeled.

### Perturbation Analysis

Through altering the dynamics of the interaction modeled, we identified specific drug targets. Perturbations upregulated or down-regulated gene expression or adjusted protein concentration. The present study incorporated 92 different perturbations on 46 constants into the baseline model to analyze their effect and possible role as therapeutic targets. For each perturbation, the constant for the respective pathways are activated or inhibited by two orders of magnitude. As all constants started with a constant value set to 1, the two possible perturbations for each constant were 100 or 0.01. Perturbations either with a magnitude fold change difference or with significant fractional differences were recorded.

## Results

Simulating the RA model and plotting through matplotlib produced Figure 3. 26 total quantities were graphed following an initial logarithmic growth pattern and then leveling out to homeostasis.

**Figure 3:**
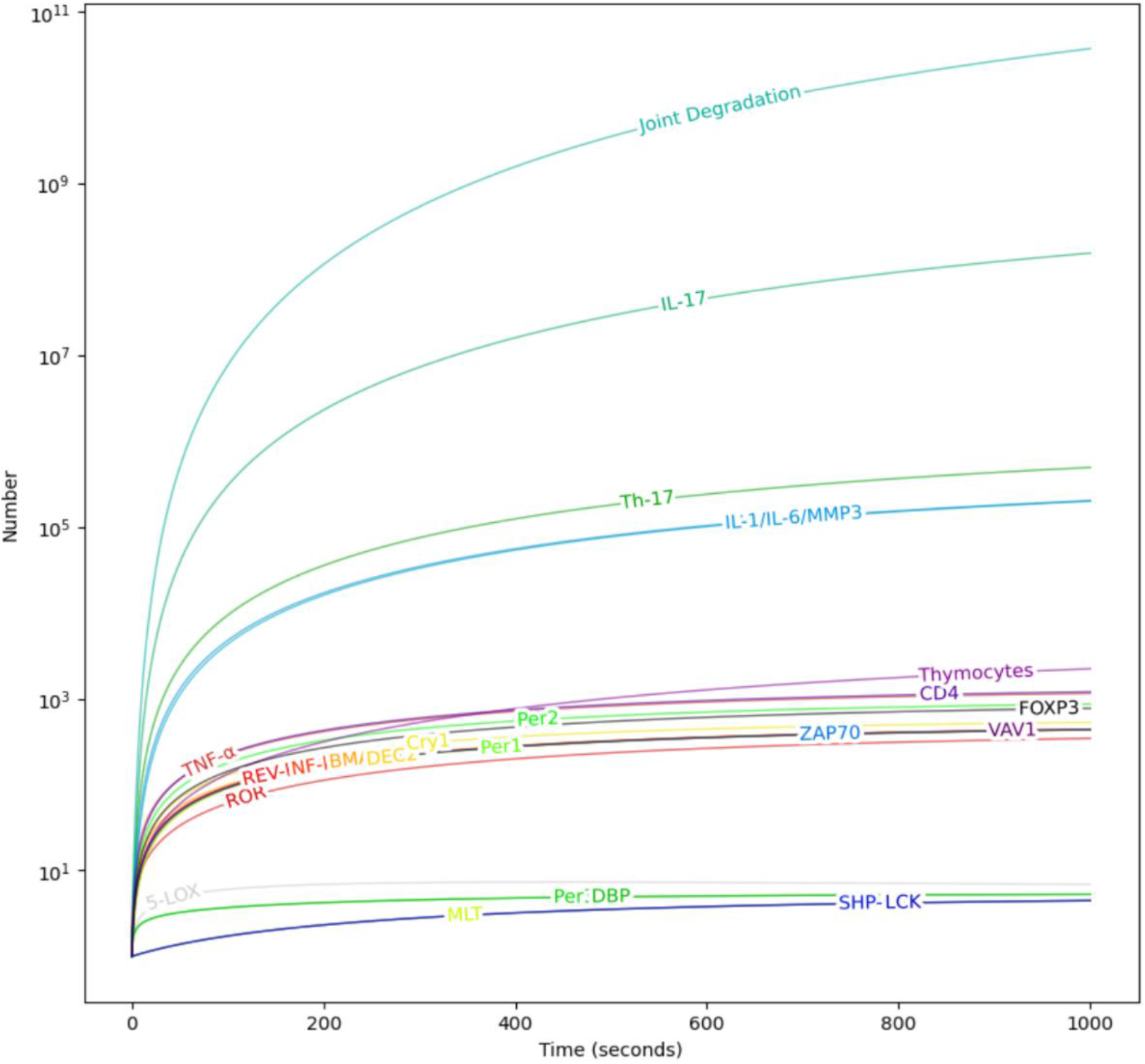
Graph of change in quantity/expression of molecules, genes, and phenomena. The x-axis represents the time in seconds and the y-axis is the count of change on a logarithmic scale. 1000 intervals of time and 10^6 intervals of counts were measured.

To quantify the change in the concentration or expression of each interactor, the following heatmap was created.

**Figure 4:**
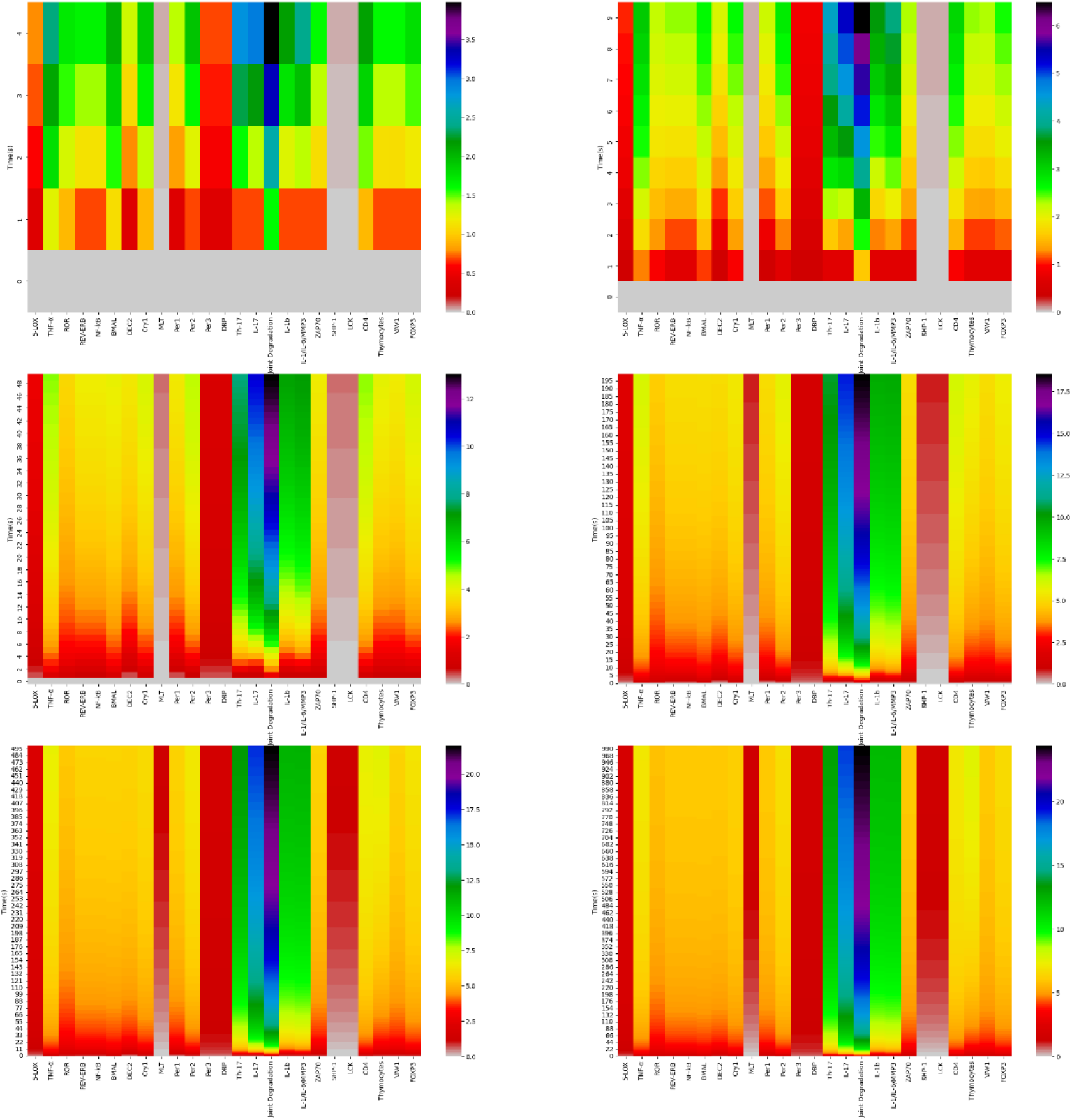
Heat maps representing the expression level or quantity of the genes, small molecules, or processes. Each graph represents a different time scale of progression in order to visualize fluctuations and expressions of concentration progression. The color progresses from red to purple, where red represents a lower level of expression/quantity and purple represents higher levels of expression/quantity.

From this heat map, we observed that after simulating the model for 5000 seconds, joint degradation was notably at a high level. Small molecules like Th-17 and IL1β had higher concentrations at t = 4950 seconds.

After simulating the baseline RA model, various perturbations were incorporated to simulate disruptions representing drugs or mutations. From testing 92 perturbations on the 46 constants, perturbations that caused significant fluctuations in joint degradation were recorded. While most perturbations caused functional changes in the same fold change range as the original joint destruction count, illustrating the ability of complex networks to maintain homeostasis, several perturbations caused a more significant difference in joint degradation. These perturbations recorded are shown in Table 4 and Table 5.

**Table 4:** Log of significant perturbations that reduced the level of joint degradation (JD) in the model. Column 2 from the left provides the reference of the constant for the respective pathway and its original value. Column 3 provides the perturbed value of the constant. Red indicates that the perturbation is upregulating the gene and green indicates that the perturbation is downregulating the gene. Column 4 presents the original value of joint destruction at time t = 4950 seconds, at which time the process had fairly reached homeostasis. Column 5 presents the perturbed value of joint destruction at time t = 4950.

| Pathway | Original Constant for Pathway | Perturbed Constant for Pathway | Change in Joint Degradation at time t = 4950 |
| --- | --- | --- | --- |
| VAV1 activating CD4 | K_22_24 = 1 | K_22_24 = 0.01 | 9.984670e+11 |
| MLT activating ROR | K_3_9 = 1 | K_3_9 = 0.01 | 7.461039e+11 |
| ROR activating Th-17 | K_14_3 = 1 | K_14_3 = 0.01 | 7.400516e+11 |
| Th-17 activating IL-17 | K_15_14 = 1 | K_15_14 = 0.01 | 1.167708e+10 |
| LCK activating CD4 | K_22_21 = 1 | K_22_21 = 0.01 | 7.606287e+11 |
| Thymocytes activating CD4 | K_22_23 = 1 | K_22_23 = 0.01 | 1.055373e+12 |
| VAV1 activating CD4 | K_22_24 = 1 | K_22_24 = 0.01 | 9.984670e+11 |
| LCK activating Thymocytes | K_23_21 = 1 | K_23_21 = 0.01 | 1.061217e+12 |

**Table 5:**
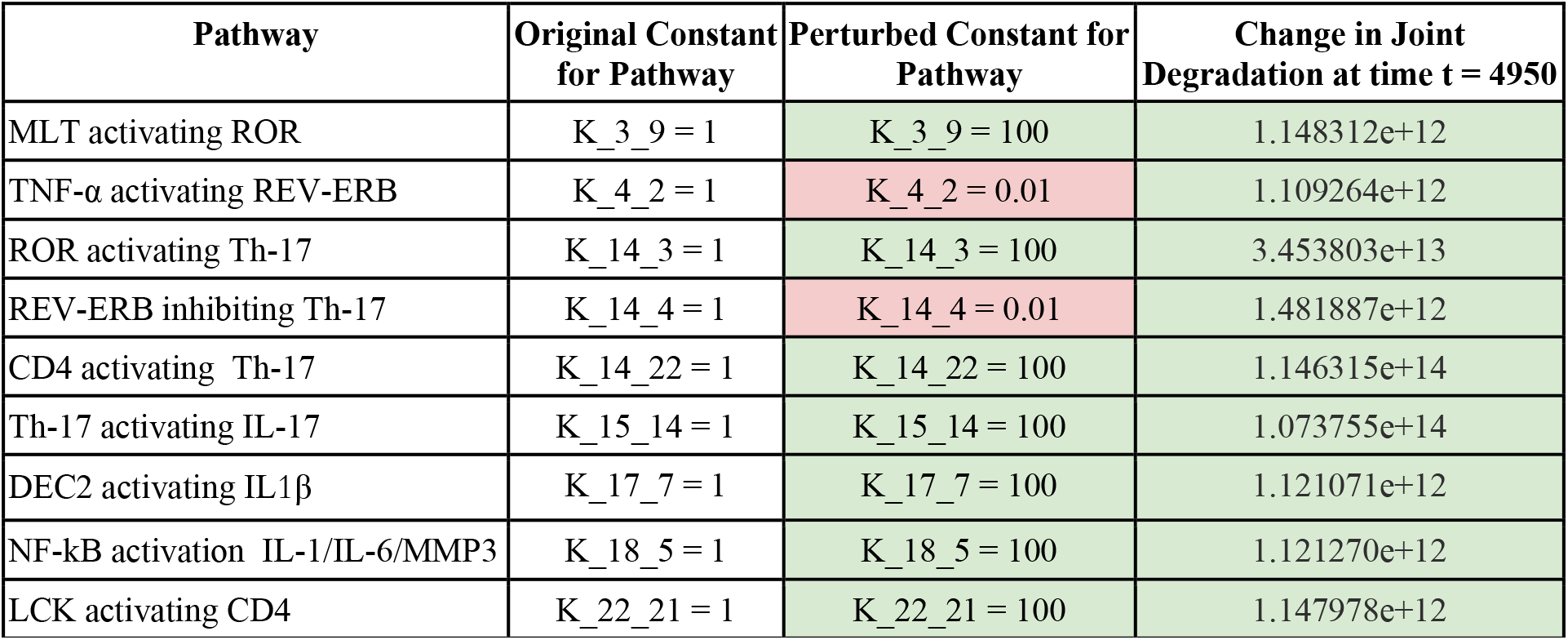
Log of significant perturbations that increased the level of joint degradation (JD). Similar table setup as in Table 4.

Notably, we found that inhibition of the VAV1-CD4, LCK-Thymocyte, MLT-ROR, ROR-Th-17, Th-17-IL-17, and LCK-CD4 pathway reduced the level of joint destruction by a fold change factor, indicating that these could potentially serve as drug targets.

Activation of the MLT-ROR, DEC2-IL1β, NF-kB-interleukin, and LCK-CD4, pathways and deregulation of the TNF-α-REV-ERB, REV-ERB-Th-17 pathways increased joint destruction, indicating that increased activity in these pathways could potentially serve as diagnostic biomarkers for higher RA risk.

## Discussion

Several pathways, when inhibited, reduced the level of joint destruction. The inhibition of these pathways could potentially serve as drug targets.

Inhibition of the LCK-CD4 pathway reduced the level of joint destruction. LCK is a gene that is known to play a role in T-cell signaling, particularly in T cell receptor (TCR) signaling. Phosphorylation by LCK of the receptor complex at the tyrosine-based activation motif (ITAM) consensus sites is found to trigger TCR signaling in patients with RA (Mellado 2015). While this relationship has been established, extensive experimentation on the potential of LCK as a novel immunosuppressant agent has not been conducted. Thus, inhibiting the specific LCK-CD4 pathway should be explored further as a potential drug target for RA.

We found that inhibition of the VAV1-CD4 pathway reduced the level of joint destruction. VAV1 is a gene that codes for the Vav1 protein which is in a family of proteins that are guanine nucleotide exchange factors (GEFs) for the Rho family GTPase and play a role in T-cell activation and development (35). Preliminary research has examined the role of a VAV1 polymorphism in RA. (36, 37). This study highlights the potential of targeting the VAV1-CD4 pathway as potential drug therapy for RA.

We found that inhibition of the MLT-ROR pathway reduced the level of joint destruction. MLT represents melatonin which is a hormone that is known to play various roles in the immune system. Particularly, Sulli et al and Cutolo et al found that varying levels of MLT are associated with various levels of pain as associated with RA as based on the human circadian rhythm (38, 39). This study highlights the importance of implementing further studies to understand the potential for regulating MLT levels to reduce pain, swelling, and other symptoms of RA. Though this is a potential avenue of drug treatment, it should be taken into consideration that regulating or adjusting levels of MLT might interfere with hormonal regulation and the circadian rhythm in the human body.

Several pathways, when activated or inhibited, increased the level of joint destruction. The exacerbated or reduced presence of these pathways could serve as diagnostic biomarkers.

Particularly, we found that activation of the DEC2-IL1β pathway increased the level of joint destruction. Previous studies found DEC2 to be increased in the synovial membrane in RA as opposed to Osteoarthritis (OA) (40). To corroborate the study and the previous findings, further experiments should be conducted to confirm the potential of DEC2 as a risk factor or diagnostic for the early identification of RA.

Activation of the NF-kB-interleukins pathway was also found to increase the level of joint destruction. Various previous studies have demonstrated the role of NF-kB in RA pathogenesis (41). While targeting NF-kB has been explored as a potential therapy for RA, this study highlights its potential as a diagnostic biomarker for RA.

Inhibition of the TNF-α-REV-ERB pathway was found to increase the level of joint destruction. TNF-α is a common target in current therapeutics for Rheumatoid Arthritis, and the relatively recent approach of targeting TNF-α has improved the success in the treatment (42). For example, Infliximab, etanercept, adalimumab, golimumab, and certolizumab-pegol are five major TNF-α inhibitor drugs that have been admitted for RA 42). While the inhibition of TNF-α has been widely explored for therapeutic purposes, there has not been much exploration into identifying lower levels of TNF-α as a diagnostic biomarker. This study highlights the potential for examining the significance of measuring TNF-α levels as a potential diagnostic criterion.

The strength of this study is that our model incorporated a wide variety of genes, proteins, and small molecules via specialized differential equations. Incorporating more components in the model allows for more accurate results. Additionally, in such a model, we can track measurements of any molecule or transcript, at any given time, allowing for the precise simulation of changes in concentration. Our model uses a well-developed basic framework and adds scientifically based proposed interactions in order to accurately simulate a particular network that has never been studied *in silico* previously. Lastly, all possible perturbations were tested in the model, allowing us to identify the most promising drug targets for RA.

Limitations of this study include several assumptions that we made. The degradation and dilution constants were derived based on assumptions of the properties of the molecules of interest. While the model assumed that that translation is happening at a slower rate than transcription such that transcription is the bottleneck, if a certain gene does not follow the assumption in a situation where translation is faster than transcription, the model would have to be altered. Additionally, 1 was used as a comparative value for the starting concentration and the constant value as it was difficult to find standardized values of concentrations and constants of specific molecules and genes in the literature. Further work includes ensuring all concentration and constants are biologically correct through running RNA sequencing to measure the number of each specific transcript in a cell at a specific time interval, running single-cell sequencing to measure a cell’s RNA transcript concentration at various intervals of time, and calculating initial concentration of a transcript without any drugs or perturbation. For proteins that were incorporated, to measure how strongly the proteins bind, binding assays should be run.

Rheumatoid Arthritis is one of the most prevalent chronic inflammatory diseases (25). Identifying more specific drug targets and diagnostic biomarkers is crucial to ensure the reduction of extensive joint destruction and bone erosion. This study highlights the role of the LCK-CD4, VAV1-CD4, and MLT-ROR pathways as potential drug targets and the role of the DEC2-IL1β, NF-kB-interleukin, and TNF-α-REV-ERB pathways as diagnostic biomarkers, and illustrates the powerful role computational modeling can play in understanding RA and other diseases.

## Acknowledgments

The authors thank the University of California, Irvine (UCI) and the Gifted and Talented Institute (GAT) for providing resources and guidance for the research.

## Supplement

The source code for the RA model can be accessed through the following link: https://colab.research.google.com/drive/1FrhARvTOewOpl7SdFdQrxon6zZYgCpZX?usp=sharing

**Table A1:** 30 interactions that were modeled between two-gene and three-gene pathways and networks. Each gene, molecule, or component was assigned a variable name that was used in the code.

| Interaction Relationship # | Molecules | Corresponding Variable Names |
| --- | --- | --- |
| 1 | 5-LOX, TNF-α, ROR | G1, G2, G3 |
| 2 | TNF-α, ROR, REV-ERB | G2, G3, G4 |
| 3 | TNF-α, ROR, NF-kB | G2, G3, G5 |
| 4 | ROR, REV-ERB, BMAL | G3, G4, G6 |
| 5 | NF-kB, BMAL, DEC2 | G5, G6, G7 |
| 6 | TNF-α, Cry1, MLT | G2, G8, G9 |
| 7 | TNF-α, Per1 | G2, G10 |
| 8 | TNF-α, Per2 | G2, G11 |
| 9 | Cry1, Per2 | G8, G11 |
| 10 | TNF-α, Per3 | G2, G12 |
| 11 | TNF-α, ROR, MLT | G2, G3, G9 |
| 12 | TNF-α, DBP | G2, G13 |
| 13 | ROR, REV-ERB, Th-17 | G3, G4, G14 |
| 14 | Th-17, IL-17, Joint Degradation | G14, G15, G16 |
| 15 | DEC2, IL1β, Joint Degradation | G7, G17, G16 |
| 16 | NF-kB, IL-1/IL-6/MMP3, Joint Destruction | G5, G18, G16 |
| 17 | ZAP70, Cry1 | G19, G8 |
| 18 | ZAP70, SHP-1 | G19, G20 |
| 19 | ZAP70, LCK | G19, G21 |
| 20 | LCK, CD4, Thymocytes | G21, G22, G23 |
| 21 | LCK, VAV1 | G21, G24 |
| 22 | VAV1, CD4 | G24, G22 |
| 23 | CD4, FOXP3 | G22, G25 |
| 24 | VAV1, SHP-1 | G24, G26 |
| 25 | CD4, Th-17 | G22, G14 |
| 26 | CD4, TNF-α | G22, G2 |
| 27 | LCK, FOXP3 | G21, G25 |
| 29 | VAV1, Joint Degradation | G24, G16 |
| 30 | FOXP3, ROR | G25, G3 |

## Notes

### Competing Interest Statement

The authors have declared no competing interest.

## References

1. Aca. “Arthritis by the Numbers: Book of Trusted Facts & Figures.” American Caregiver Association, 17 Nov. 2018, americancaregiverassociation.org/2018/11/arthritis-by-the-numbers-book-of-trusted-facts-figures/

2. “Arthritis and RA.” Global RA Network, globalranetwork.org/project/disease-info/.

3. How Rheumatoid Arthritis Affects More Than Joints, www.arthritis.org/diseases/more-about/how-rheumatoid-arthritis-affects-more-than-joints.

4. Young, A. “Which Patients Stop Working Because of Rheumatoid Arthritis? Results of Five Years’ Follow up in 732 Patients from the Early RA Study (ERAS).” Annals of the Rheumatic Diseases, vol. 61, no. 4, 2002, pp. 335–340., doi:10.1136/ard.61.4.335.

5. “Rheumatoid Arthritis (RA).” Centers for Disease Control and Prevention, Centers for Disease Control and Prevention, 27 July 2020, www.cdc.gov/arthritis/basics/rheumatoid-arthritis.html.

6. “Rheumatoid Arthritis.” Mayo Clinic, Mayo Foundation for Medical Education and Research, 18 May 2021, www.mayoclinic.org/diseases-conditions/rheumatoid-arthritis/diagnosis-treatment/drc-20353653#:~:text=There is no cure for, modifying antirheumatic drugs (DMARDs).

7. Lewis, Myles J., et al. “Molecular Portraits of Early Rheumatoid Arthritis Identify Clinical and Treatment Response Phenotypes.” Cell Reports, vol. 28, no. 9, 2019, doi:10.1016/j.celrep.2019.07.091.

8. “If Your RA Biologic Doesn’t Help Enough: What Next?” WebMD, WebMD, www.webmd.com/rheumatoid-arthritis/first-biologic-fails.

9. “Methotrexate (Oral).” Methotrexate (Oral) | Michigan Medicine, www.uofmhealth.org/health-library/d00060a1.

10. Emami, Jaber, and Zahra Ansarypour. “Receptor Targeting Drug Delivery Strategies and Prospects in the Treatment of Rheumatoid Arthritis.” Research in Pharmaceutical Sciences, vol. 14, no. 6, 2019, p. 471., doi:10.4103/1735-5362.272534.

11. Ostrowska, Monika, et al. “Cartilage and Bone Damage in Rheumatoid Arthritis.” Reumatologia/Rheumatology, vol. 56, no. 2, 2018, pp. 111–120., doi:10.5114/reum.2018.75523.

12. Smolen, Josef S, et al. “Tocilizumab Inhibits Progression of Joint Damage in Rheumatoid Arthritis Irrespective of Its Anti-Inflammatory Effects: Disassociation of the Link between Inflammation and Destruction.”.” Annals of the Rheumatic Diseases, vol. 71, no. 5, 2011, pp. 687–693., doi:10.1136/annrheumdis-2011-200395.

13. Yap, Hooi-Yeen, et al. “Pathogenic Role of Immune Cells in Rheumatoid Arthritis: Implications in Clinical Treatment and Biomarker Development.” Cells, vol. 7, no. 10, 2018, p. 161., doi:10.3390/cells7100161.

14. “Biomarker Tests for Rheumatoid Arthritis Improve Care.” Rheumatoid Arthritis, 24 Nov. 2015, blog.arthritis.org/rheumatoid-arthritis/biomarker-tests-rheumatoid-arthritis/.

15. Tomlin, Claire J., and Jeffrey D. Axelrod. “Biology by Numbers: Mathematical Modelling in Developmental Biology.” Nature Reviews Genetics, vol. 8, no. 5, 2007, pp. 331–340., doi:10.1038/nrg2098.

16. “Mathematical Models in Biology - WebHome Kuttler/ · Chapter 1 Introduction to Modelling Literature:” Dokumen.tips, dokumen.tips/documents/mathematical-models-in-biology-webhome-kuttler-chapter-1-introduction-to.html.

17. Daun, Silvia, et al. “Equation-Based Models of Dynamic Biological Systems.” Journal of Critical Care, vol. 23, no. 4, 2008, pp. 585–594., doi:10.1016/j.jcrc.2008.02.003.

18. Google Colaboratory, Google, colab.research.google.com/notebooks/intro.ipynb?utm_source=scs-index.

19. Covert, Markus W.,Fundamentals of Systems Biology: from Synthetic Circuits to Whole-Cell Models. CRC Press, 2017.

20. Liu, Enze, et al. “Gene Regulatory Network Review.” Encyclopedia of Bioinformatics and Computational Biology, 2019, pp. 155–164., doi:10.1016/b978-0-12-809633-8.20218-5.

21. Philips, Ron Milo & Ron. “” How Fast Do RNAs and Proteins Degrade?” Cell Biology by the Numbers How Fast Do RNAs and Proteins Degrade Comments, book.bionumbers.org/how-fast-do-rnas-and-proteins-degrade/.

22. Jahanban-Esfahlan, Rana, et al. “Melatonin in Regulation of Inflammatory Pathways in Rheumatoid Arthritis and Osteoarthritis: Involvement of Circadian Clock Genes.” British Journal of Pharmacology, vol. 175, no. 16, 2017, pp. 3230–3238., doi:10.1111/bph.13898.

23. Chang, Christina, et al. “The Nuclear Receptor REV-ERBα Modulates Th17 Cell-Mediated Autoimmune Disease.” Proceedings of the National Academy of Sciences, vol. 116, no. 37, 2019, pp. 18528–18536., doi:10.1073/pnas.1907563116.

24. Amir, Mohammed, et al. “REV-ERBα Regulates TH17 Cell Development and Autoimmunity.” Cell Reports, vol. 25, no. 13, 2018, doi:10.1016/j.celrep.2018.11.101.

25. Robert, Marie, and Pierre Miossec. “IL-17 in Rheumatoid Arthritis and Precision Medicine: From Synovitis Expression to Circulating Bioactive Levels.” Frontiers in Medicine, Google Colaboratory, Google, colab.research.google.com/notebooks/intro.ipynb?utm_source=scs-index.

26. Nakahira, Kumiko, et al. “Phosphorylation of FOXP3 by LCK Downregulates MMP9 Expression and Represses Cell Invasion.” PLoS ONE, vol. 8, no. 10, 2013, doi:10.1371/journal.pone.0077099.

27. Li, Zhiyuan, et al. “FOXP3 Regulatory T Cells and Their Functional Regulation.” Cellular & Molecular Immunology, vol. 12, no. 5, 2015, pp. 558–565., doi:10.1038/cmi.2015.10.

28. Han, J, et al. “Lck Regulates Vav Activation of Members of the Rho Family of GTPases.” Molecular and Cellular Biology, vol. 17, no. 3, 1997, pp. 1346–1353., doi:10.1128/mcb.17.3.1346.

29. Hehner, Steffen P., et al. “Tyrosine-Phosphorylated Vav1 as a Point of Integration for T-Cell Receptor- and CD28-Mediated Activation of JNK, p38, and Interleukin-2 Transcription.” Journal of Biological Chemistry, vol. 275, no. 24, 2000, pp. 18160–18171., doi:10.1074/jbc.275.24.18160.

30. Kassem, Sahar, et al. “A Natural Variant of the T Cell Receptor-Signaling Molecule Vav1 Reduces Both Effector T Cell Functions and Susceptibility to Neuroinflammation.” PLOS Genetics, vol. 12, no. 7, 2016, doi:10.1371/journal.pgen.1006185.

31. Lewintre, E. J., et al. “Cryptochrome-1 Expression: a New Prognostic Marker in B-Cell Chronic Lymphocytic Leukemia.” Haematologica, vol. 94, no. 2, 2009, pp. 280–284., doi:10.3324/haematol.13052.

32. Habashy, Deena Mohamed, et al. “Cryptochrome-1 Gene Expression Is a Reliable Prognostic Indicator in Egyptian Patients with Chronic Lymphocytic Leukemia: A Cohort Prospective Study.” Turkish Journal of Hematology, 2017, doi:10.4274/tjh.2017.0169.

33. Castro-Sánchez, Patricia, and Pedro Roda-Navarro. “Physiology and Pathology of Autoimmune Diseases: Role of CD4 T Cells in Rheumatoid Arthritis.” Physiology and Pathology of Immunology, 2017, doi:10.5772/intechopen.70239.

34. Tesmer, Laura A., et al. “Th17 Cells in Human Disease.” Immunological Reviews, vol. 223, no. 1, 2008, pp. 87–113., doi:10.1111/j.1600-065x.2008.00628.x.

35. Tybulewicz, Victor L. J., et al. “Vav1: a Key Signal Transducer Downstream of the TCR.” Immunological Reviews, vol. 192, no. 1, 2003, pp. 42–52., doi:10.1034/j.1600-065x.2003.00032.x.

36. Latawa, Paridhi. “Identification and Analysis of Novel Synovial Tissue-Based Biomarkers and Interacting Pathways for Rheumatoid Arthritis.” 2021, doi:10.1101/2021.03.19.21253995.

37. “Novel Small Molecule Inhibitor of Rheumatoid Arthritis.” Novel Small Molecule Inhibitor of Rheumatoid Arthritis | SBIR.gov, www.sbir.gov/sbirsearch/detail/76336.

38. Sulli, A., et al. “Melatonin Serum Levels in Rheumatoid Arthritis.” Annals of the New York Academy of Sciences, vol. 966, no. 1, 2002, pp. 276–283., doi:10.1111/j.1749-6632.2002.tb04227.x.

39. Cutolo, M. “Circadian Rhythms: Glucocorticoids and Arthritis.” Annals of the New York Academy of Sciences, vol. 1069, no. 1, 2006, pp. 289–299., doi:10.1196/annals.1351.027.

40. Olkkonen, Juri, et al. “Differentially Expressed in Chondrocytes 2 (DEC2) Increases the Expression of IL-1β and Is Abundantly Present in Synovial Membrane in Rheumatoid Arthritis.” Plos One, vol. 10, no. 12, 2015, doi:10.1371/journal.pone.0145279.

41. Samimi, Leila Nejatbakhsh, et al. “NF-KB Signaling in Rheumatoid Arthritis with Focus on Fibroblast-like Synoviocytes.” Autoimmunity Highlights, vol. 11, no. 1, 2020, doi:10.1186/s13317-020-00135-z.

42. Radner, Helga, and Daniel Aletaha. “Anti-TNF in Rheumatoid Arthritis: an Overview.” Wiener Medizinische Wochenschrift, vol. 165, no. 1-2, 2015, pp. 3–9., doi:10.1007/s10354-015-0344-y.

